# Cell-type and cytosine context-specific evolution of DNA methylation in the human brain

**DOI:** 10.1101/2020.07.14.203034

**Authors:** Hyeonsoo Jeong, Isabel Mendizabal, Stefano Berto, Paramita Chatterjee, Thomas Layman, Noriyoshi Usui, Kazuya Toriumi, Connor Douglas, Devika Singh, Iksoo Huh, Todd M. Preuss, Genevieve Konopka, Soojin V. Yi

**Affiliations:** School of Biological Sciences, Georgia Institute of Technology, Atlanta, GA 30332, USA; Center for Cooperative Research in Biosciences (CIC bioGUNE), Basque Research and Technology Alliance (BRTA), Bizkaia Technology Park, Building 801A, Derio, 48160, Spain; Department of Neuroscience, UT Southwestern Medical Center, Dallas, TX 75390, USA; Center for Medical Research and Education and Department of Neuroscience and Cell Biology, Graduate School of Medicine, Osaka University, Suita, Osaka 565-0871, Japan; Schizophrenia Research Project, Department of Psychiatry and Behavioral Sciences, Tokyo Metropolitan Institute of Medical Science, Tokyo, 156-8506, Japan; College of Nursing and The Research Institute of Nursing Science, Seoul National University, Seoul, 03080, South Korea; Division of Neuropharmacology and Neurologic Diseases, Yerkes National Primate Research Center, Emory University, and Department of Pathology, Emory University School of Medicine, Atlanta, GA 30329, USA

## Abstract

Cell-type specific epigenetic modifications are critical for brain development and neuropsychiatric diseases. Here we elucidate evolutionary origins of neuron- and oligodendrocyte-specific DNA methylation in human prefrontal cortex, and demonstrate dynamic and distinctive changes of CG and CH methylation. We show that the human brain has experienced pronounced reduction of CG methylation during evolution, which significantly contributed to cell-type specific active regulatory regions. On the other hand, a substantial increase of CH methylation occurred during human brain evolution, associated with fine-tuning expression in development and neuronal subtypes. The majority of differential CG methylation between neurons and oligodendrocytes originated before the divergence of hominoids and catarrhine monkeys, and carries strong signal for genetic risk for schizophrenia. Remarkably, a substantial portion of differential CG methylation between neurons and oligodendrocytes emerged in the human lineage and harbors additional genetic risk for schizophrenia, implicating epigenetic evolution of human cortex in increased vulnerability to neuropsychiatric diseases.

## INTRODUCTION

DNA methylation is a stable epigenetic modification of genomic DNA with critical roles in brain development ^1,2^. Comparisons of the whole-genome DNA methylomes of humans and chimpanzees suggested a reduction of human brain specific CG methylation associated with gene regulation ^3,4^. Methylation at non-CG contexts (CH methylation, where H = A, C, T) is also present in human and mouse brains, where it is associated with postnatal neuronal maturation and cell-type specific transcriptional activity ^1,5,6^. Cell-type specific epigenetic marks, including DNA methylation and histone modifications, underlie cell-type specific gene expression and disease susceptibility in humans ^7,8^. To understand the contribution of DNA methylation to human brain-specific gene regulation and disease susceptibility, it is necessary to extend our knowledge of evolutionary changes in DNA methylation during human brain evolution. Here, we present novel comparative analyses of neuron- and oligodendrocyte-specific whole-genome DNA methylomes of humans, chimpanzees, and rhesus macaques. By integrating these data with transcriptome data from the same individuals ^9^ and recent data from studies of bulk and cell-type specific epigenetic and transcriptomic modifications of human brains ^2,10–12^, we demonstrate dramatic changes of DNA methylation in a cell-type and cytosine context-specific manner during human brain evolution, impacting the human brain regulatory landscape and disease susceptibility.

## RESULTS

### Distinctive methylomes of neurons and oligodendrocytes in human and non-human primate prefrontal cortex

We generated cell-type specific DNA methylomes of sorted nuclei from post-mortem brain samples of humans ^8^, chimpanzees (*Pan troglodytes*), and rhesus macaques (*Macaca mulatta*). We selected Brodmann area 46 (BA46) from dorsolateral prefrontal cortex (also referred to as ‘prefrontal cortex’ or ‘cortex’ henceforth), which is involved in higher-order cognitive functions that have likely undergone marked changes in human evolution ^13,14^. Neuronal (NeuN+) and oligodendrocyte (OLIG2+) cell populations were isolated using fluorescence-activated nuclei sorting (FANS) as previously described ^8,9^. We used wholegenome bisulfite sequencing (WGBS) to generate DNA methylomes at nucleotide resolution for NeuN+ and OLIG2+ populations (Supplementary Figure S1). Altogether, we compared 25, 11, 15 NeuN+ methylomes and 20, 11, 13 OLIG2+ methylomes from human, chimpanzee, and rhesus macaque, respectively (Supplementary Tables S1 and S2). We also performed whole-genome sequencing (WGS) of the same individuals (Supplementary Table S3). The mean coverages for the WGBS and WGS data are 20.6X (±8.8) and 23.2X (±5.9), respectively. Polymorphic sites at cytosines (i.e. C to T for forward strand and G to A for reverse strand) were excluded to avoid spurious methylation calls due to the technical limitation of distinguishing bisulfite-converted thymine from unmethylated cytosine (Supplementary Table S4).

As in humans, non-human primate prefrontal cortex is highly methylated at CG sites, and NeuN+ DNA is more highly methylated than OLIG2+ DNA (*P* < 10^−100^, two-sample K-S tests, Figure 1A). In comparison, CH methylation occurs in much lower frequencies than CG methylation, and is nearly exclusive to NeuN+ DNA in humans ^1,8^ and non-human primates (Figure 1A). CH methylation is higher in humans compared to chimpanzees (Figure 1A, P = 0.03, Mann-Whitney U test using proportions of mCH > 10%), and humans and chimpanzees are more highly CH methylated than rhesus macaques and mice (Figure 1A, P = 4.3 x 10^−5^, Kruskal-Wallis test). Principal component analyses demonstrate that cell type explains the largest amount of variation in both methylation contexts, followed by species (Figure 1B). Since OLIG2+ DNA generally lacks CH methylation, there is little separation of species for CH OLIG2+ (Figure 1B). As the genomic patterns and cellular distributions of CG and CH methylation are highly distinct from each other (Figure 1B), we analyzed them separately.

**Figure 1.**
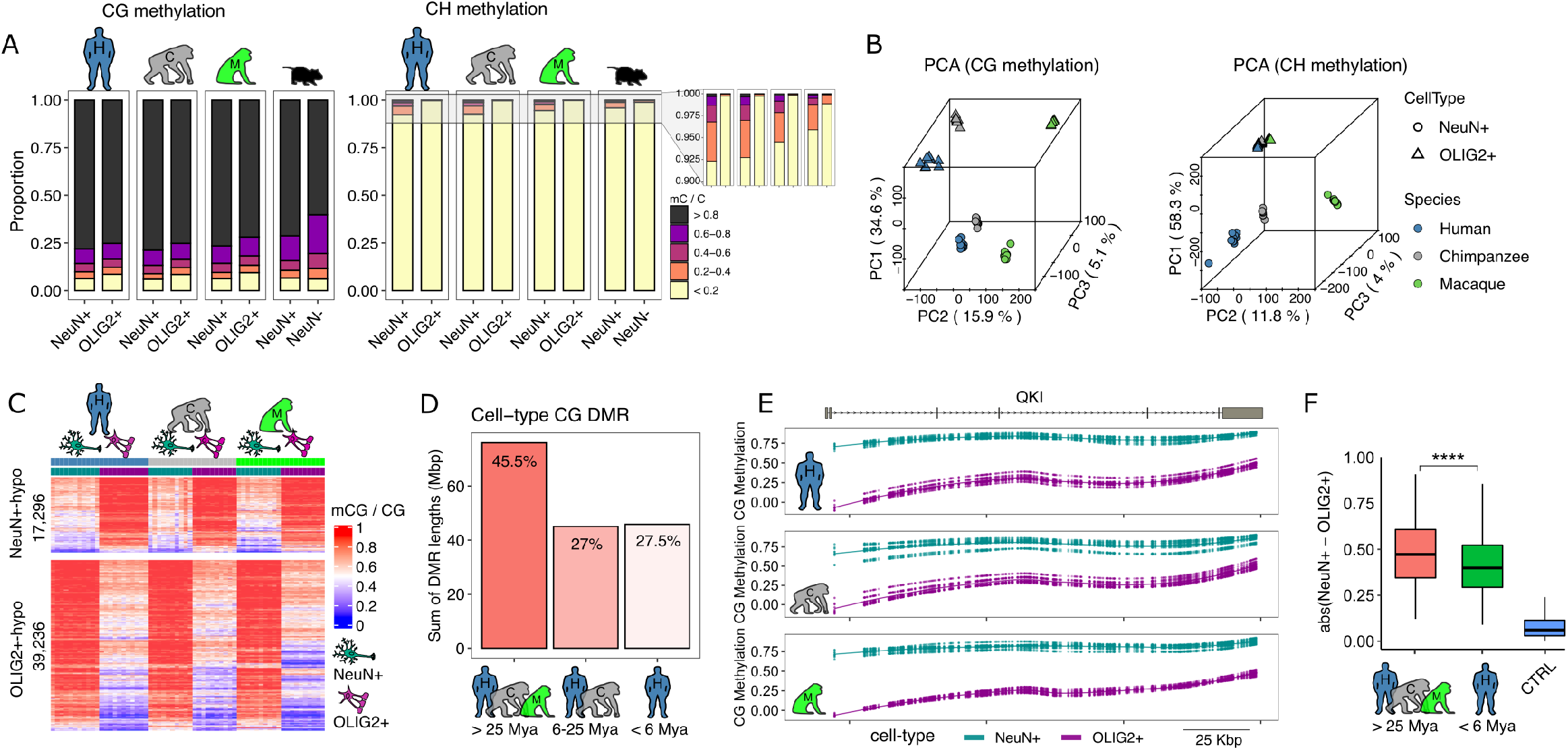
CG and CH methylation in neurons (NeuN+ cells) and oligodendrocytes (OLIG2+ cells) in human and non-human primate prefrontal cortex. A) The proportions of methylated CG and CH sites. Human and non-human primate neurons and oligodendrocytes are highly CG methylated. Human and non-human primate neurons show low levels of CH methylation and oligodendrocytes show even lower levels. CH methylation is highest in human neurons, followed by chimpanzees, rhesus macaques, and mice. B) Principal component analysis of methylated cytosines in two contexts (CG and CH). The top two principal components (PCs), PC1 and PC2, distinguish cell type and species, respectively. C) CG methylation levels in neurons (left columns for each species) and oligodendrocytes (right columns for each species). A greater number of DMRs are hypermethylated in neurons (red, in the left columns) compared to oligodendrocytes (right columns). D) Approximately half (45.5%) of CG DMRs differentially methylated between NeuN+ and OLIG2+ cells are conserved in all three species, with 27% conserved between humans and chimpanzees, and 27.5% specific to the human. E) An example CG DMR between NeuN+ and OLIG+ cells spanning the entirety of *QKI*, a gene involved in oligodendrocyte function. This gene is consistently hypomethylated in oligodendrocytes compared to neurons in all three species. F) The absolute methylation difference of NeuN+ and OLIG2+ cells is highest for DMRs conserved in all three species compared to those specific to the human. Methylation difference between NeuN+ and OLIG2+ cells calculated from genomic regions serving as statistical control (CTRL), with matched number of CG and G+C nucleotide contents, are also displayed. Asterisks represent P-value < 0.00001 (Mann-Whitney U-test).

### Conservation and divergence of cell-type-specific CG methylation

Due to the high rate of CG mutations associated with DNA methylation ^15^ only 9.6 million CG sites (out of 28 million total human CGs) are conserved in all three species (Supplementary Figure S2), and these sites are biased toward hypomethylation (Supplementary Figure S3). To avoid bias associated with CG conservation, we first identified differentially methylated regions (DMRs) that distinguish humans and chimpanzees using conserved sites (21 out of 25 million CGs analyzed), and subsequently added DNA methylation data from rhesus macaques to polarize direction of evolutionary change (Methods). In this analysis, we applied methods developed for the analysis of whole genome bisulfite sequencing data to identify species, cell-type and interaction effects on DNA methylation while taking into account variation due to sex, age, and bisulfite conversion rates (Methods).

Non-human primate methylomes of NeuN+ and OLIG2+ are highly distinct from each other and show clear clustering of cell types in each species (Supplementary Figure S4), as in humans ^8^. There are 56,532 CG DMRs (75.9 Mbp) between NeuN+ and OLIG2+ DNA that are conserved in all three species (Figures 1C and Supplementary Table S5). These conserved DMRs account for nearly 50% of all DMRs between NeuN+ and OLIG2+ in humans (Figure 1D). Consequently, a large portion of differential CG methylation between NeuN+ and OLIG2+ DNA originated before the divergence of hominoids and catarrhine monkeys. Enrichment tests utilizing cis-regulatory interactions based on long-range regulatory domains ^16^ show that these regions are highly enriched in genes harboring functions specific to neurons and oligodendrocytes (Supplementary Table S6). For example, we show one conserved DMR spanning the whole *QKI* locus (Figure 1E). This gene, which is an RNA binding protein involved in myelination and oligodendrocyte differentiation ^17^, is encompassed in its entirety by a DMR in all three species so that it is hypomethylated in oligodendrocytes while hypermethylated in neurons. Gene expression data from matched samples ^9^ shows that *QKI* is up-regulated in oligodendrocytes compared to neurons in all three species (P < 10^−7^ in all three species, Methods), indicating that cell-type specific regulation may be facilitated by DNA methylation. Interestingly, the absolute methylation difference between neurons and oligodendrocytes was significantly more pronounced in the evolutionarily ‘old’ DMRs conserved in all three species compared to those recently evolved in human (Figure 1F; results of the same comparison using chimpanzee DMRs are also consistent, Supplementary Figure S5).

### Pronounced CG hypomethylation of human prefrontal cortex and human neuron-specific regulatory landscape

We found 23,703 CG DMRs (13.1Mbp) that experienced differential CG methylation since the divergence of humans and chimpanzees (Methods, Figures 2A and 2B), distributed across different functional regions, including regions currently annotated as non-coding intergenic regions (Supplementary Figure S6). These CG DMRs include 7,861 for which both cell types are differentially methylated between humans and chimpanzees (4,253 human-specific and 3,608 chimpanzee-specific CG DMRs, based on the comparison to macaques). The rest of the CG DMRs show DNA methylation changes in a cell-type-specific manner in each species (Figures 2A and Supplementary Table S7). In the following we present results of analyses for each cell type, combining DMRs that are common in both cell types and DMRs that are cell-type-specific in each species (Methods).

**Figure 2.**
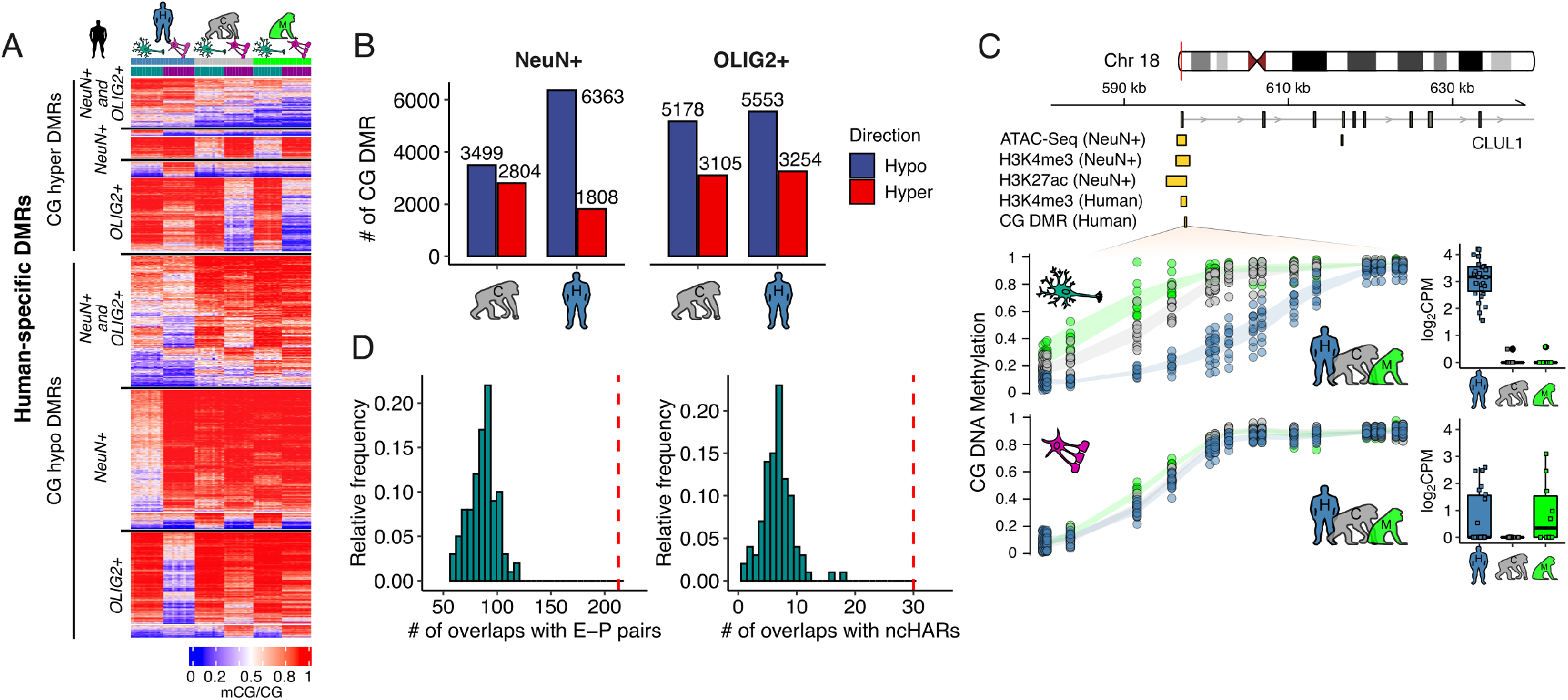
Evolutionary changes in CG methylation. A) Heatmap representation of mean DNA methylation of all 23,703 human DMRs in the three species illustrates dramatic reduction of CG methylation in human prefrontal cortex, especially in neurons. B) Numbers of DMRs in NeuN+ and OLIG2+ cells in human and chimpanzee frontal cortex. C) An example of the relationship between human neuron-hypo CG DMR and other epigenetic marks in the *CLUL1* locus, a gene widely expressed in the brain. This DMR overlaps with multiple other epigenetic marks of active chromatin in the human brain, including neuron-specific ATAC-Seq peak, neuron-specific H3K4me3 peak, neuron-specific H3K27ac peak. This DMR also overlaps with a human-specific brain H3K4me3 peak compared to chimpanzee and macaque. D) Human neuron-hypo CG DMRs are significantly co-localized with enhancer-promoter (E-P) pairs and ncHARs (red dashed lines). Null distributions were plotted based on the GC matched control region sets (n=100) that overlap with enhancerpromoter pairs and ncHARs. Chimpanzee neuron-hypo CG DMRs did not show such patterns (Supplementary Figure S9).

CG DMRs tend to show reduction of DNA methylation (hypomethylation) in human prefrontal cortex compared to chimpanzee in both cell-types (Figure 2B). While most previous studies focused on neurons, recent studies have begun to unveil the functional and evolutionary importance of oligodendrocytes-specific changes ^9,18^. Indeed, we identified a substantial number of human-derived hypomethylated DMRs specific to oligodendrocytes (Figure 2A and 2B). To examine the transcriptional impact of cell-type epigenetic specializations of the human brain, we analyzed gene expression data from the same individuals and nuclei populations ^9^. Hypomethylation near transcription start sites is associated with up-regulation of genes ^19^. Consistent with this, genes harboring CG DMRs are significantly enriched in differentially expressed genes between humans and chimpanzees in oligodendrocytes and neurons (P = 0.002 and P = 0.01 for NeuN+ and OLIG2+, respectively, hypergeometric test). In addition, species differences in DNA methylation present significant negative correlations with expression differences (Supplementary Figure S10). These results indicate widespread and significant contributions of evolutionarily recent DNA methylation changes in human oligodendrocytes to the transcriptional landscape of the human brain.

Human neurons in particular harbor a large number of hypomethylated CG DMRs compared to chimpanzee neurons (Figure 2B, 6,363 hypomethylated CG DMRs in human neurons versus 3,499 hypomethylated DMRs in chimpanzee neurons, OR = 2.82, P = 5.5 x 10^−20^, chisquare test). Taking advantage of recent functional genomics data from human neurons, we show that human neuron-specific hypomethylated CG DMRs (referred to as ‘neuron-hypo CG DMRs’ henceforth) mark active regulatory regions of the neuronal genome (Supplementary Figure S7). Specifically, a substantial portion of human neuron-hypo CG DMRs co-localize with brain-specific enhancers (Supplementary Figure S8), as well as other recently characterized cell-type specific human brain epigenetic marks, including neuron-specific H3K27ac (fold-enrichment = 3.1, P < 0.01, permutation test), H3K4me3 (foldenrichment = 8.5, P < 0.01), and ATAC-Seq (fold-enrichment = 8.2, P < 0.01) peaks ^7,20^ For example, we show a neuron-hypo CG DMR in a 5’ region of the *CLUL1* locus, which overlaps with other epigenomic signatures of active chromatin marks observed in human neurons (Figure 2C). This gene is broadly expressed in brain, although its functional role in the brain function is not resolved yet. We also report human brain and human neuron-specific up-regulation from matched gene expression data (Figure 2C), indicating that CG hypomethylation and other epigenetic modifications that have occurred in the human prefrontal cortex since the divergence of humans and chimpanzees are implicated in human brain specific up-regulation of this gene.

In order to reveal the target genes of these epigenetically coordinated regulatory elements in human neurons, we integrated three-dimensional maps of chromatin contacts from the developing human cortex ^21^. This analysis identified 213 enhancer-promoter pairs (Figure 2D, fold-enrichment = 2.45, P < 0.01, permutation test), supporting physical chromatin interactions between spatially adjacent human neuron-hypo CG DMRs in human neuron nuclei (Supplementary Table S8). Interestingly, genes affected by these enhancer-promoter interactions are enriched in functional categories including neuron differentiation and development (Supplementary Table S9). We also explored the co-occurrence of epigenetically identified regulatory elements with those emerging from DNA sequence analyses. Non-coding human accelerated regions (ncHAR) significantly overlap with human-specific hypomethylated CG DMRs (Figure 2D, fold-enrichment = 4.45, P < 0.01, permutation test). In contrast, chimpanzee-specific hypomethylated CG DMRs did not show significant patterns (Supplementary Figure S9). Notably, ncHARs also show an excess of three-dimensional interactions with distant human hypo CG DMRs, which include 7 experimentally validated human brain enhancer ncHARs ^22^. In addition, human neuro-hypo CG DMRs frequently co-occur with human neuron-specific histone H3-trimethyl-lysine 4 (H3K4me3) modification ^11^ (fold-enrichment = 18.1, empirical P-value < 0.01, permutation test). Taken together, these results demonstrate the confluence of human-derived genetic and epigenetic innovations, and that CG hypomethylation of human neurons contributed to the active chromatin landscape of human prefrontal cortex in a cell-type specific manner.

### Signature of evolutionarily recent CH hypermethylation in human neurons

CH methylation is limited to a few cell types in the body ^23,24^, and occurs at much lower frequency than CG methylation (Figure 1A). Nucleotide substitution rates at CH sites and CH methylation do not have a significant correlation ^25^. Consequently, we were able to follow the evolutionary dynamics of CH methylation for the majority of CH positions. Among the 1.1 billion CH positions examined in the human genome, 716 million sites (71.2%) were found in the three species we examined (Methods). We found 51.9 million CH sites hypermethylated in NeuN+ compared to OLIG2+ DNA (FDR < 0.05). Among these, 23.6 million sites (45.5%) show NeuN+ DNA hypermethylation in all three species. Human and chimpanzee neurons share an additional 16.3 million (31.4%) CH hypermethylated sites not found in macaque (Figure 3A). Moreover, an additional 3.1 million CH sites gained methylation in the human neurons (Supplementary Figure S11), which is a significant excess compared to the 2.2 million sites gained via CH methylation in the chimpanzee neurons (OR = 1.54, 95% CI 1.534 – 1.546, P < 10^−20^, chi-square test). Thus, in contrast to the pronounced hypomethylation in the CG context, human neurons are predominantly hypermethylated (Figure 3B) compared to other primates.

**Figure 3.**
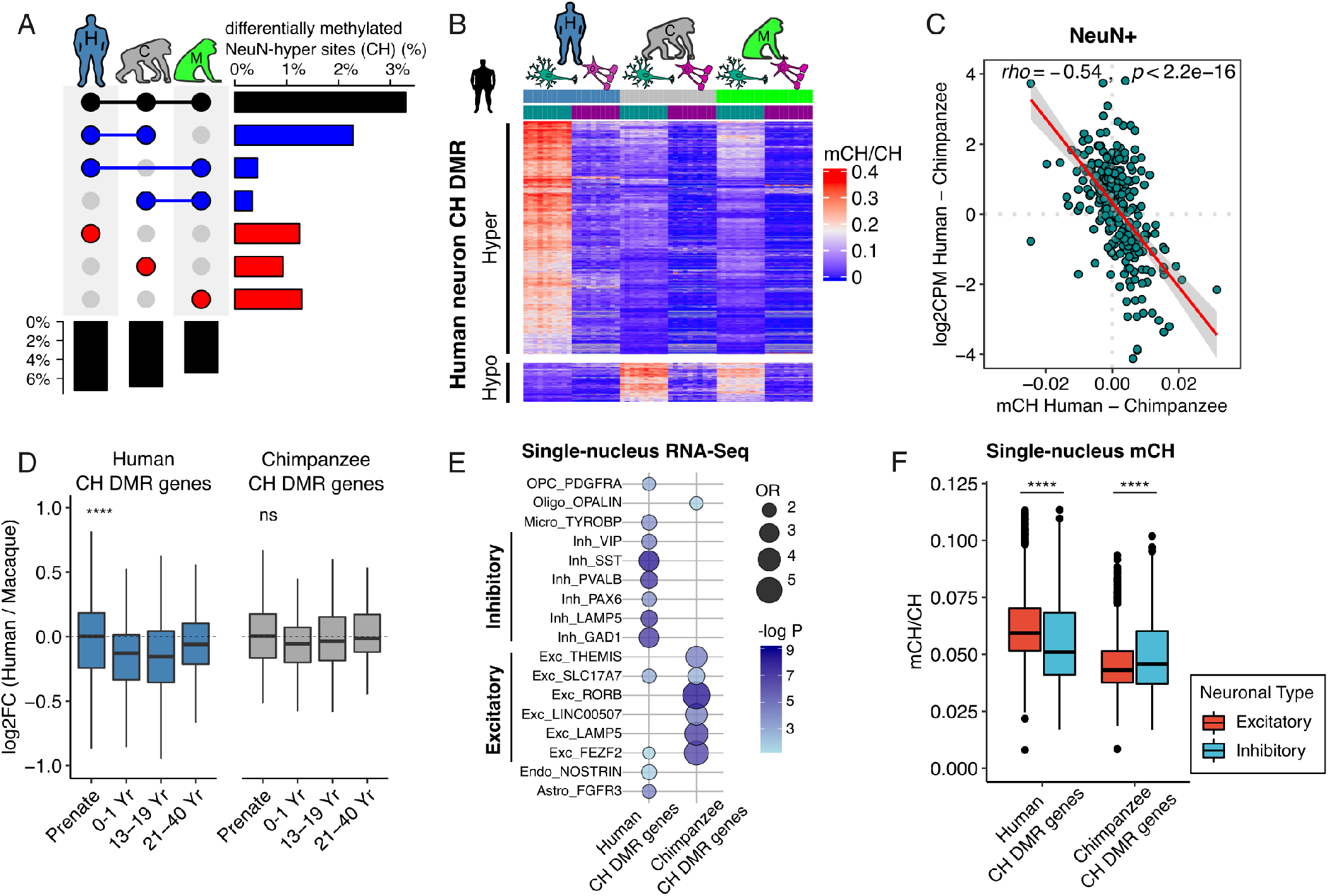
Evolutionary changes in CH methylation. CH hypermethylation is significantly higher in human neurons compared to other primates. A) Differences in the proportions of sites with neuronal CH methylation between species. B) Mean methylation levels of human-specific CH DMRs demonstrate pronounced hypermethylation of human neurons. C) CH methylation between humans and chimpanzees strongly predicts gene expression difference. D) Gene expression fold-change between human and macaque in CH DMR genes across developmental time points. Macaque samples were age-matched to human developmental time points in a previous study ^12^. Asterisks represent P-value < 0.00001 (Kruskal-Wallis test). E) Enrichment of human and chimpanzee CH DMR genes in specific cell-types. Human CH DMR genes are enriched in inhibitory neurons whereas chimpanzee CH DMR genes are enriched in excitatory neurons. In each gene set, genes expressed in at least 50% of the cells that are statistically significant (FDR < 0.05 and log_2_FC > 0.3) are included. Celltype data are from human medial temporal gyrus (MTG) ^27^. OR = Odds Ratio. F) CH methylation of neuronal subtypes for CH DMR genes using methylation of single nuclei from the human frontal cortex ^10^. Human CH DMR genes are hypomethylated in inhibitory neurons whereas chimpanzee CH DMR genes are hypomethylated in excitatory neurons. Asterisks represent P-value < 0.00001 (Mann-Whitney U-test).

CH methylation of gene bodies is one of the strongest predictors of repression of gene expression in humans and mice ^1,5,8,26^. We find similarly strong repressive effects of genic CH methylation on gene expression in human and non-human primate neurons (Supplementary Figures S12). Moreover, differential CH methylation between species is strongly negatively correlated with gene-expression differences between species, indicating that the change of CH methylation is a major determinant of neuronal transcriptional divergence (Figure 3C). CH methylation is nearly absent in fetal brains and accumulates rapidly after birth ^1^. We thus hypothesized that the repressive impact of CH methylation might be more pronounced in early postnatal development, and subsequently examined gene expression data from bulk brain tissue during development ^12^. Indeed, genes bearing signatures of human-specific CH methylation accumulation (referred to as human CH DMR genes, Supplementary Table S10, Methods) are similarly expressed in human and macaque brains during prenatal growth but show reduced expression in humans following birth (Figure 3D). In contrast, chimpanzee CH DMR genes do not exhibit such a pattern (Figure 3D and Supplementary Figure S13). We integrated our data with those from sorted neurons from individuals of different ages ^2,9^, to examine cell-type differences. Human CH DMR-genes showed lower expression in neurons than in non-neurons or oligodendrocytes in most developmental stages, and the reduction of neuronal expression was more evident in toddler and early teen data compared to data from adults (Supplementary Figure S14).

Interestingly, human CH DMR genes are significantly enriched in gene sets representing inhibitory neurons, based on single-nucleus transcriptome data from the middle temporal gyrus ^27^ (Figure 3E), as well as those previously identified as markers of inhibitory neurons ^10,28,29^ (FE = 5.3, P < 0.0001, permutation test). Moreover, these genes were more highly methylated in excitatory neurons than in inhibitory neurons in single-nucleus DNA methylation data from the same brain region ^10^ (Figure 3F and Supplementary Figure S15). Integrating these observations, we hypothesize that human-specific CH methylation of inhibitory-neuron-specific genes may silence their expression in the genomes of excitatory neurons, thereby promoting functional specificity of neuron subtypes. On the other hand, chimpanzee CH DMR genes, which represent regions with no evolutionary change in humans, are enriched in human excitatory neuron markers (Figure 3E) and are slightly, albeit significantly, more highly methylated in inhibitory neurons (Figure 3F).

### Human neuron-specific CG methylation contributes additional risk to schizophrenia heritability

We have previously shown that genomic regions exhibiting differential CG methylation between neurons and oligodendrocytes are associated with increased risk for neuropsychiatric disorders, especially for schizophrenia ^8^. Other studies have noted that sites of differential histone modification ^7^ or DNA methylation ^26^ between neurons and nonneurons significantly contribute to heritability for neuropsychiatric disorders. Our data can provide further insights into the evolution of genetic risk for neuropsychiatric disorders.

We used the stratified linkage disequilibrium score regression framework ^30^ to estimate the contribution of DMRs to the genetic heritability of various diseases and complex traits. Since evolutionary DMRs are generally extremely short (e.g. the median lengths of human CG DMR and CH DMR are 471 bps and 246 bps, respectively) we extended the DMR windows by 25kbp on both sides to improve the confidence intervals of the estimates (Methods). To avoid potential bias, we removed the MHC region from this analysis as in other studies ^30^. We found a strong enrichment of risk for schizophrenia and other brain-related traits at neuron-hypo CG DMRs that are evolutionarily conserved in the three species, while no signal was detected at OLIG2+ conserved DMRs (Figure 4A and Supplementary Tables S11 and S12). Non-brain traits such as height, which is extremely polygenic, and body mass index (BMI) were also detected, consistent with the previously proposed role of the central nervous system in the genetic architecture of BMI ^30^. Moreover, human-specific neuron-hypo CG DMRs exhibited significant enrichment for schizophrenia heritability (Figure 4A). In contrast, chimpanzee neuron-hypo CG DMRs did not show significant enrichment for any human trait. A sliding-windows analysis further demonstrates that the aforementioned signal for schizophrenia was centered at the DMRs and did not originate from extended adjacent regions (Figure 4B). In contrast, both conserved and human-specific CH DMRs show significant depletion for schizophrenia heritability (Figures 4A and 4B). Notably, the depletion signal was centered around the CH DMRs, whereas no other diseases (with the exception of bipolar disorder) nor chimpanzee-specific regions showed a significant trend (Figure 4A), implying that CH hypermethylated genomic regions are devoid of common DNA polymorphisms associated specifically with schizophrenia.

**Figure 4.**
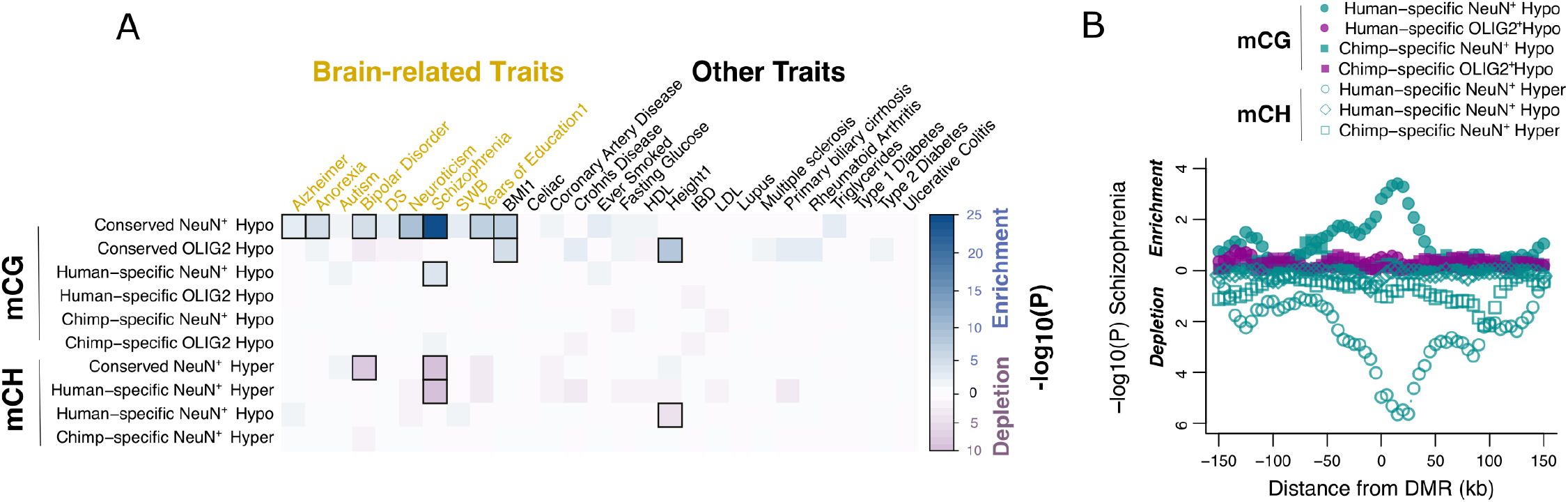
Evolutionarily derived DMRs contribute to brain disease susceptibility. A) Significance levels for the enrichment for genetic heritability in different DMRs (+/−25kb) and complex traits. Both conserved and human-specific neuronal DMRs are associated with schizophrenia. Enrichment with FDR < 0.05 are highlighted in squares. Notably, CG DMRs hypomethylated in NeuN+ cells compared to OLIG2+ cells in all three species (conserved NeuN+ hypo) are highly enriched in variants for several brain-related traits, and human-specific NeuN+ hypo shows enrichment in schizophrenia. B) Sliding window analysis of enrichment for schizophrenia in DMRs. The Y-axis represents the P-values in sliding windows around DMRs classified by species (human or chimpanzee), cell-type (NeuN+ or OLIG2+), and cytosine context (mCG or mCH).

## DISCUSSION

Previous whole-genome methylome analyses of the human brain have demonstrated that differential DNA methylation plays critical roles in brain development ^1,31^, cell-type differentiation ^1,8^, and disease susceptibility ^8,26^. Despite such importance for genome regulation and human health, the ways DNA methylation and other epigenetic mechanisms have changed during human brain evolution have not previously been characterized at the cell-type level. Reliable identification of human-specific epigenetic modifications at the celltype level has been a limiting factor in previous studies due to the heterogeneity of brain tissue and the different relative cell compositions of different species. Our cell-type specific whole-genome methylomes of neurons and oligodendrocytes from humans, chimpanzees, and rhesus macaques enable us, for the first time, to elucidate evolutionary changes of DNA methylation during human brain evolution. We have previously discovered an excess of CG hypomethylation in human prefrontal cortex compared to chimpanzee ^3^, and many of these sites are located in noncoding regulatory regions of the human genome ^4^. We find this to be the case for both neurons and oligodendrocytes, which could contribute to increased gene expression levels that have been reported in human brains ^9,32,33^. Intriguingly, CH methylation is higher in human and chimpanzee prefrontal cortex compared to rhesus macaque and mice. Also, we show that human prefrontal cortex has higher CH methylation than chimpanzees and rhesus macaques. Therefore, our evidence suggests that CH methylation in the prefrontal cortex increased during the evolution of primates. However, data from brain tissue of a wider variety primates and other mammals are needed to rigorously test this hypothesis. CH methylation is highly negatively correlated with gene expression and is a strong predictor of gene-expression divergence between species. Integrating our results with developmental bulk tissue data and single-cell functional genomics data from human brains, we show that the human-specific increase of CH methylation might be particularly important for early human brain development and finetuning neuron subtype cell identities. Additional data from matched brain regions are needed to investigate evolutionary changes of CH methylation in relation to cell-type specificity. Due to the limitation of bisulfite sequencing, our data cannot separate methylcytosines from hydroxymethylcytosines (hmCs), which might play distinctive roles in neuron subtypes ^34^. While additional data are needed, currently available maps ^35^ do not suggest a significant impact of hmC on the differential methylation patterns identified in this study (Supplementary Figure S17).

Epigenetic differences between neurons and non-neuronal cells are prevalent in non-coding regions and locate in regions that account for schizophrenia heritability ^7,8,26^. Here we show that genomic regions with differential CG methylation between neurons and oligodendrocytes that contribute greatest to schizophrenia risk originated before the emergence of the catarrhine ancestor. Moreover, CG positions that underwent hypomethylation since the divergence of humans and chimpanzees, despite their relatively small proportion, provide additional significant genetic risk to schizophrenia. These results provide a link between human brain evolution and heritability for schizophrenia. Therefore, our analyses confirm that evolutionary changes in cell-type-specific alteration of DNA methylation and other epigenetic mechanisms play an important role in the expression of genes characteristic of human brains and shape our genetic risk to neuropsychiatric disorders.

## Acknowledgements

This work was partially supported by the Asan Foundation (Biomedical Science Scholarship) to H.J; Uehara Memorial Foundation to N.U.; JSPS Grant-in-Aid for Early-Career Scientists (18K14814) to N.U. and Scientific Research (C) (18K06977) to K.T.; Takeda Science Foundation to N.U.; the JSPS Program for Advancing Strategic International Networks to Accelerate the Circulation of Talented Researchers (S2603) to S.B., N.U., K.T. and G.K.; the James S. McDonnell Foundation 21^st^ Century Science Initiative in Understanding Human Cognition – Scholar Award and the Jon Heighten Scholar in Autism Research at UT Southwestern to G.K.; National Science Foundation (SBE-131719) to S.V.Y; the National Chimpanzee Brain Resource, NIH R24NS092988, and the NIMH (MH103517), to T.M.P., G.K., and S.V.Y.

## Materials and Methods

### Sample acquisition, whole-genome sequencing and whole-genome bisulfite sequencing

Information on samples used in this work was previously described in Berto et al. ^36^. Briefly, adult human postmortem brain samples from Brodmann area 46 (BA46) were acquired from the National Institutes of Health NeurobioBank (the Harvard Brain Tissue Resource Center, the Human Brain and Spinal Fluid Resource Center, VA West Los Angeles Healthcare Center, and the University of Miami Brain Endowment Bank). These samples included 25 and 22 NeuN+ and OLIG2+ specimens, respectively. All non-human primate samples were obtained from homologous regions in chimpanzees (NeuN+ *n* = 11, OLIG2+ n = 11) and rhesus macaques (NeuN+ *n* = 15, OLIG2+ *n* = 13). These are the same tissue samples as used in Berto et al. ^36^.

### Whole-genome bisulfite data processing

We followed the same data processing steps described in our previous work ^37^. Briefly, the methylome libraries were diluted and loaded onto Illumina HiSeqX system for sequencing using 150bp paired-end reads. We performed quality and adapter trimming using TrimGalore v.0.4.1 (Babraham Institute) with default parameters. Reads were mapped first to PhiX genome (NC_001422.1) to remove the spike-in control and the remaining reads were subsequently mapped to the chimpanzee PanTro5 and macaque rheMac8 reference genomes using Bismark v 0.14.5 ^38^ and bowtie v2.3.4 ^39^. After de-duplication, we obtained coverage for over 84% of the CpGs in the chimpanzee genome with an average read depth 19.32x, and over 91% of CpGs in the macaque genome with an average read depth of 21.61x. We calculated fractional methylation (ratio of the number of methylated cytosine reads to the total number of reads) levels at individual cytosines. Bisulfite conversion rates were estimated by mapping the reads to the lambda phage genome (NC_001416.1).

### Whole-genome sequencing data processing

Quality and adapter trimming was performed using TrimGalore v.0.4.1 (Babraham Institute) with default parameters. Reads were mapped to the hg19, PanTro5 or rheMac8 reference genomes using BWA v0.7.4 ^40^ and duplicates were removed using picard v2.8.3 (https://broadinstitute.github.io/picard/index.html). We identified genetic polymorphisms from re-sequencing data following the GATK v4 best practices workflow ^41^. For base recalibration, we used vcf files for known variants from dbSNP for chimpanzee and macaque from the following links: ftp://ftp.ncbi.nlm.nih.gov/snp/organisms/chimpanzee_9598/VCF/ and ftp://ftp.ncbi.nlm.nih.gov/snp/organisms/macaque_9544/VCF/. We applied hard filters for genotype calling with the following parameters: --filterExpression “QD < 2.0 || FS > 60.0 || MQ < 40.0 || MQRankSum < −12.5 || ReadPosRankSum < −8.0”. For chimpanzee, we identified 10,980,856 variants with mean depth >24x. For macaque, we identified 30,001,119 variants with mean depth >24x. Since C>T and G>A polymorphisms at CpG sites can generate spurious differential methylation patterns, we removed polymorphic CpGs from downstream differential methylation analyses keeping a total of 26,024,877 and 24,740,404 non-polymorphic CpGs for chimpanzee and macaque genomes, respectively. For quality control of SNP calling, we performed principal component analyses using additional chimpanzee and bonobo samples from de Manuel et al. ^42^ using 75,575 common SNPs from chromosome 20. As expected, our chimpanzee samples clustered with other chimpanzees and not with bonobos (Supplementary Figure S18). We recapitulated the genetic ancestry of de Manuel et al. samples and identified most of our individuals as Western chimpanzees (*Pan troglodytes verus*) while one sample (sample ID Anja) clustered with Nigeria-Cameroon chimpanzees (*Pan troglodytes ellioti*).

### RNA-Seq data

We used our previously generated matched samples of RNA-Seq datasets for human (without brain-related diseases), chimpanzee, and rhesus macaques from GSE108066, GSE107638, and GSE123936.

### Liftover of non-human primates cytosine positions to human genome

We lifted over the non-human primates’ cytosine coordinates to human hg19 genome using UCSC batch liftover tool (panTro5ToHg19.over.chain.gz and rheMac8ToHg19.over.chain.gz for chimpanzee and rhesus macaque, respectively). For the CG DMR analysis, we did not perform three-way species analyses based on lifted over coordinates due to the rapid evolutionary loss of CG sites since the macaque split. Compared to around 21 million CG sites conserved between human and chimpanzee, only around 9.6 million CGs are conserved between human and macaque, whereas 13 million CGs in macaque show non-CG dinucleotides in human. To circumvent this issue, we first identified human-chimpanzee differentially methylated regions (DMRs) using conserved CGs and then used orthologue regions in the macaque rheMac8 genome to polarize the DMRs (see *“Incorporation of Rhesus Macaque as an outgroup species”* for additional details). We removed cytosines located in paralogous sequences in at least one species to avoid erroneous mapping (i.e. one-to-many or many-to-one mapping between species). For the CH methylation analysis, we used orthologous cytosines conserved among the three species.

### Identification of CG differential methylation

We identified differentially methylated positions of 1) cell-types (NeuN+ vs. OLIG2+), 2) species (human vs. chimpanzee where both cell types show the same direction and magnitude of methylation differences between two species) and 3) cell-type-specific species changes (either cell-type exclusively shows DNA methylation difference between species) using DSS (ver. 2.3) Bioconductor package ^43^. DSS handles variance across biological replicates and models read counts from WGBS experiments while accounting for additional biological factors. Specifically, we considered age (converted to three level categorical variable), sex, and conversion rates as covariates in the model.

Fractional methylation ~ cell_type + species + species:cell_type + sex + age_class + conversion_rates

To remove low coverage loci, we only included sites with at least 5x coverage in 80% of individuals per species or cell-type. We used a false discovery rate (FDR) threshold of 5% to identify significant differentially methylated positions. For DMR identification, we considered a minimum length of 50bp with at least 4 significant differentially methylated positions. We removed cell-type DMRs and species DMRs that overlap with cell-type-specific species changes (i.e. interaction of cell-type and species effects) to remove redundant DMRs. We only considered the DMRs that show >10% of average methylation difference between human and chimpanzee for species DMR and >15% of average methylation difference between cell-types for cell-type DMR (please also see the section *“Incorporation of Rhesus Macaque as an outgroup species”* for detailed explanation of final set of DMRs).

Of note, as our differential methylation analyses were run under a multifactor design in DSS, the estimated coefficients in the regression were based on a generalized linear model framework using the arcsine link function to reduce dependence of variance on the fractional methylation levels ^44^. The distribution of the statistic is determined by the differences in methylation levels as well as by biological and technical factors such as read depth. The sign of the test statistic indicates the direction of methylation. However, the values of the test statistic cannot be directly interpreted as fractional methylation differences. For DMRs, the tool generates “areaStat” values which are defined as the sum of the test statistic of all CG sites within the DMR. To identify the stringent sets of DMRs we excluded DMRs if the average test statistics of corresponding CGs in the region (areaStat divided by the number of CGs) was below the test statistic corresponding to FDR = 0.05.

### Incorporation of rhesus macaque as an outgroup species

We retrieved the corresponding genomic coordinates in rheMac8 using the Ensembl Primate EPO multiple sequence alignment ^45^. Read counts and methylation values of the CGs in corresponding regions were obtained from the macaque samples. Only CG sites with at least 5x coverage in 80% of the individuals per species were considered. The DMRs resulting from human and chimpanzee samples that had low alignment coverages with macaque (<50%) or included less than 4 CGs in macaque were considered “unclassified” DMRs. After adding macaque data, we fitted a beta regression model using the average methylation level of each individual accounting for the covariates indicated above. Among the cell-type DMRs resulting from human and chimpanzee samples, DMRs in which macaque showed cell-type changes in the same direction and exhibited >15% fractional methylation difference were considered conserved cell-type DMRs.

We then used stringent criteria to categorize the species specificity of DMRs as human- or chimpanzee-specific. For example, a human-specific hypomethylated DMR should satisfy the following criteria: 1) the average fractional methylation of human is significantly lower than that of chimpanzee and macaque (FDR < 0.05), 2) the absolute methylation difference between human and macaque is greater than that between chimpanzee and macaque, 3) the proportion of the absolute methylation difference between human and macaque is greater than 5%, and 4) both of the two cell-types satisfy these criteria. Those DMRs that did not satisfy these criteria were considered “unclassified”. We used the same logic to specify human-specific hypermethylated DMRs and chimpanzee-specific hypo- and hypermethylated DMRs.

We also examined species-specific DMRs which show differential methylation between species but exclusively in one cell-type (i.e. either cell-type shows differential methylation patterns derived from either the human or chimpanzee lineage).

### Identification of CH differential methylation

Unlike CG methylation, >70% of cytosine positions were conserved among the three species. Thus, we used orthologous cytosines across the three species to infer differentially methylated positions. Because CH methylation is sensitive to bisulfite conversion rate ^46^, we only used individuals with high bisulfite conversion rates (>99.5%). We down-sampled and matched sample size across the species to avoid any bias derived from the different sample sizes across groups (N=11 for each species and cell-type). We removed sites in which >50% of individuals in at least one group have fewer than 5 read counts.

For each CH site, we fitted a generalized linear model using the arcsine function to identify differentially methylated CH positions among species adjusting for other covariates (age, sex, and bisulfite conversion rate) using DSS. To fit our parsimonious approach, we also performed pair-wise analyses between species considering all combinations (i.e. human vs. chimpanzee, human vs. macaque, and chimpanzee vs. macaque). Benjamini–Hochberg correction (FDR) was used to perform multiple comparisons. We used the parsimonious approach to detect species-specific methylation changes with a cutoff of fractional methylation difference between species > 10% and FDR < 0.05. For example, human-specific CH methylated sites showed FDR < 0.05 from both human vs. chimpanzee and human vs. macaque comparisons and FDR > 0.05 from the chimpanzee vs. macaque comparison as well as a >10% difference of fractional methylation in humans compared to both chimpanzee and macaque fractional methylation levels.

To identify human-specific and chimpanzee-specific CH DMRs, we identified significantly differentially methylated regions between human and chimpanzee using the differentially methylated positions generated from a human-chimpanzee comparison. We considered a minimum region of 50bp with at least 4 significant differentially methylated positions (FDR < 0.05) and covering >10 cytosines. Similarly, we used an average methylation difference of 10% as a cutoff. Using average methylation of macaque from corresponding regions, we detected human-specific and chimpanzee-specific CH DMRs using the following criteria. Human-specific CH DMRs are defined as DMRs that show a significant human-chimp difference with at least 4 differentially methylated positions as well as a methylation difference between human and macaque of >5% that is also greater than the methylation difference between chimpanzee and macaque. Similarly, chimp-specific CH DMRs are DMRs that satisfy the following criteria: a significant human-chimp difference with at least 4 differentially methylated positions and a methylation difference between chimpanzee and macaque of >5% that is also greater than methylation difference between human and macaque. To obtain regions in which both human and chimpanzee were differentially methylated compared to macaque, we checked the overlap between human-macaque CH DMRs and chimpanzee-macaque CH DMRs.

### Identification of DMR genes

To identify differentially methylated genes, we extracted genes with at least one DMR within a 3kb window upstream and downstream of the gene body. To remove redundant genes among different categories of DMR genes, we used average gene body methylation as an additional indicator to assign genes into the DMR gene category using the following criteria. Human-specific hyper CH DMR-genes are defined as DMR genes that include at least one human-specific hyper CH DMR and show higher average gene body methylation compared to the average gene body methylation of chimpanzee and macaque. Also, the absolute methylation difference between human and macaque should be greater than the methylation difference between chimpanzee and macaque.

### CH methylation of neuronal subtypes

We examined methylation patterns of neuronal subtypes for CH DMR genes. Average gene body methylation of CH DMR genes was calculated for neuronal cells from 21 human neuronal subtypes ^47^. For the marker gene analysis of neuron subtypes, we used known excitatory and inhibitory neuron markers from Luo et al. 2017 ^47^. We included the marker genes that are orthologous to the three species. These include 20 excitatory neuron markers (*SATB2, TYRO3, ARPP21, SLC17A7, TBR1, CAMK2A, ITPKA, ABI2, RASAL1, FOXP1, SLC8A2, SV2B, PTPRD, LTK, LINGO1, NRGN, NPAS4, KCNH3, BAIAP2, ARPP19*) and 13 inhibitory neuron markers (*ERBB4, GAD1, SLC6A1, CCNE1, EPHB6, KCNAB3, LPP, TBC1D9, DUSP10, KCNMB2, UBASH3B, MAF, ANK1*).

### Lineage-specific accelerated non-coding regions

We used a set of human accelerated regions from Capra et al. ^48^, which combined regions identified from independent studies (i.e. the 721 ‘Pollard HARs’ from Lindblad-Toh et al. ^49^, the 1356 ‘ANC’ regions from Bird et al. ^50^, the 992 ‘HACNS’ regions from Prabhakar et al. ^51^, and the 63 ‘Bush08’ regions from Bush and Lahn ^52^). Statistical significance and foldenrichment for DMRs were computed from the occurrences of DMRs for each feature compared to GC matched control region sets (n=100).

### Hydroxymethylation

We used previously published methylome and hydroxymethylome maps at nucleotide resolution in the adult human brain ^53^. The hmC and mC sites were defined in the original paper. We included the cytosines that are orthologous across the three species (*n* = 2,905,389). We compared the proportions of differentially methylated loci between 5-hydroxymethylcytosines (hmC) and 5-methylcytosines (mC). The proportions of the differentially methylated loci at hmC loci (4.2%) and mC loci (4.2%) showed no difference.

### Contribution of DMRs to disease heritability using stratified LD score regression

To quantify the contribution of DMRs to the genetic risk of different traits and diseases, we performed stratified LD score regression analyses ^54,55^. This method estimates the percentage of heritability explained by a set of SNPs in a certain trait using GWAS summary statistics and computes the enrichment and significance by comparing the observed heritability to the expectation given the fraction of the genome considered. We used default parameters and excluded the MHC region as in Finucane et al. 2015. Together with the DMR annotations, we also included the basal functional categories described in the original paper. The list of GWAS traits and references are listed in Supplementary Table 11.

The stratified LD score regression method produces large standard errors when the annotation categories cover a small fraction of the genome. The total cumulative length of the DMRs in our study, especially the species-derived ones, were considerably smaller than those used in Finucane et al. ^54^. Therefore, we extended the DMR regions 25kb on both sides. To ensure the GWAS signals were centered around the DMRs and not emerging from the extended regions, we ran stratified LD score regression in sliding windows 300kb around the DMRs with a window size of 20kb and step size of 5kb.

Conserved CG DMRs were more numerous and longer than human-specific ones, which could lead to increased statistical power on stratified LD score regression analyses. In order to directly compare the significance of conserved and human-specific DMR categories to schizophrenia heritability, we performed partitioned stratified LD score analyses using 100 random sub-samplings of conserved regions. As shown in Supplementary Figure 15A-B, even when comparable datasets were used, conserved neuron hypomethylated DMRs consistently showed larger enrichments than human-derived ones. Similarly, subsampling of human hyper CH DMR to chimpanzee hyper DMR number and length showed stronger depletion in the former (Supplementary Figure 15 C-D). These analyses indicate that the observed differential patterns are not due to different DMR number and lengths.

## Notes

### Competing Interest Statement

The authors have declared no competing interest.

### Summary of Updates

updated the acknowledgement section.

